# Evolution of anisogamy in the early diverging fungus, *Allomyces*

**DOI:** 10.1101/230292

**Authors:** Sujal S. Phadke, Shawn M. Rupp, Melissa A. Wilson Sayres

## Abstract

Gamete size dimorphism between sexes (anisogamy) is predicted to have evolved from an isogamous system in which sexes have equal-sized, monomorphic gametes. Although adaptive explanations for the evolution of anisogamy abound, we lack comparable insights into molecular changes that bring about the transition from monomorphism to dimorphism. The basal fungal clade *Allomyces* provides unique opportunities to investigate genomic changes that are associated with this transition in closely related species that show either isogamous or anisogamous mating systems. The anisogamous species show sexual dimorphism in gamete size, number, pigmentation and motility. We sequenced transcriptomes of five *Allomyces* isolates representing the two mating systems, including both male and female phenotypes in the anisogamous species. Maximum likelihood ancestral character state reconstruction performed in MESQUITE using the de-novo assembled transcriptomes indicated that anisogamy likely evolved once in *Allomyces*, and is a derived character as predicted in theory. We found that sexual stages of *Allomyces* express homologs of several genes known to be involved in sex determination in model organisms including *Drosophila* and humans. Furthermore, expression of *CatSper* homologs in male- and female-biased samples in our analysis support the hypothesis that gamete interaction in the anisogamous species of *Allomyces* may involve similar molecular events as the egg-sperm interaction in animals, including humans. Although the strains representing either mating system shared much of the transcriptome, supporting recent common ancestry, the analysis of rate of evolution using individual gene trees indicates high substitution rates and divergence between the strains. In summary, we find that anisogamy likely evolved once in *Allomyces*, using convergent mechanisms to those in other taxa.

## Introduction

Anisogamy is a mating system in which gamete size varies between sexes (Bell 1978). The ubiquity of anisogamy across eukaryotes has motivated numerous theoretical studies into its adaptive significance (Parker, Baker, and Smith 1972, Parker 1982, Murlas Cosmides and Tooby 1981). These studies propose that four selective pressures including disruptive selection, sperm competition, sperm limitation, and cytoplasmic conflicts, together or alone, can explain the evolution of anisogamy (Togashi and Cox 2011). Empirical support for each selective mechanism has been found in organisms as diverse as microscopic green algae (Togashi et al. 2012, Kuroiwa 2010; Miyamura 2010; Greiner, Sobanski, and Bock 2015) and various animal species with internal (Gage and Morrow 2003) or external fertilization (Levitan 1993).

Although the proposed selective pressures vary, all theoretical studies agree that anisogamy likely evolved from an ancestral isogamous mating system in which gametes of the two sexes were of equal size (Matsuda and Abrams 1999; Togashi and Cox 2011). Studies in marine algae have corroborated this theoretical prediction. Specifically, Nozaki et al (2006) showed that the male-specific gene in anisogamous algae *Pledorina starii* is a homolog of the gene specifying one of the mating types in the related isogamous species *Chlamydomonas reinhardtii*. This indicated that maleness in volvocean algae was probably established from the isogametic mating type during the evolution of anisogamy (Nozaki et al. 2006). Multiple studies have verified this genetic continuity between isogamous and anisogamous algal lineages by demonstrating how evolution has reprogrammed homologs of this *(MID)* gene to control sexual phenotype in anisogamous species (Ferris et al. 2010; Nozaki 2008; Nozaki et al. 2014). However, the isogamous and anisogamous systems in the algal lineages are taxonomically widespread and genetically and ecologically highly divergent, preventing direct comparison of mating systems at the genetic level to isolate the causes and consequences of anisogamy. Because there are few other biological systems known to provide such comparative context, molecular studies on anisogamy are largely restricted to algae. We are far from understanding the genetic basis of transitions to anisogamy in other eukaryotes in which anisogamy likely evolved independently (Togashi and Cox 2011; Togashi et al. 2012; Mignerot and Coelho 2016).

The fungal genus *Allomyces* is one of the few experimentally tractable models that shows a continuum of isogamy and anisogamy in closely related species, providing an ideal system with which to analyze genetic changes associated with the transition to anisogamy. *Allomyces* is a genus of filamentous fungi in the phylum Blastocladiomycota, which emerges at the base of Ophisthokonts (the clade shared by animals and fungi) in the eukaryotic tree of life (James et al. 2006). Three groups of species have been identified within *Allomyces* based on the morphological details of lifecycle stages, *Eu-Allomyces* species with anisogamous mating systems, *Cystogenes* with isogamous systems and *Brachy-Allomyces* containing species without any known form of sexual reproduction (Emerson 1941; Olson 1984). Porter et al (2011) found such morpho-species based classification to be inconsistent with the rDNA based phylogeny in which isolates that were previously assigned to the same ‘morphospecies’ occupy distinct clades (Porter et al. 2011). This raised the possibility of polyphyletic origin and frequent transitions between mating systems within *Allomyces*, a hypothesis that could be tested using a phylogenetic analysis of additional isolates and loci.

With the exception of the model species, *A. macrogynus*, there are no genomic resources for *Allomyces*. In contrast, the sexual cycles are relatively well characterized. The anisogamous species, *A. macrogynus* and *A. arbusculus*, are hermaphrodites that form two types of gamete-sacs called gametangia (Morrison 1977). One is a colorless gametangium, which releases relatively large, colorless gametes compared to the orange gametangia that release smaller, orange colored gametes. Keeping with common convention, we refer to the larger colorless gametes as females and the smaller, pigmented gametes as males. Both gamete types are motile, although females are slower than males (Pommerville 1978). The mechanism of gamete interaction has not been fully elucidated; however, the female gametes are known to secrete a pheromone called sirenin, which attracts conspecific and heterospecific male gametes (Pommerville 1977; Machlis 1968). Recently, Syeda et al (2016) showed that synthetic sirenin can mimic the action of mammalian hormone, progesterone, and induce hyperactivation by causing Ca^2+^ influx through the *CatSper* channels in human sperm (Syeda et al. 2016). Homologs of genes encoding *CatSper* channels have been found in the genome of the model anisogamous species, *A. macrogynus* (Cai and Clapham 2012); however, it is not known whether they are involved in sexual interactions. Yet, there is some evidence that the interaction between male and female gametes of the anisogamous species depends on Ca^2^+ ions (Pommerville, Strickland, and Harding 1990). These studies suggest that the mechanisms of gamete interaction through *CatSper* channels may be conserved, or convergently evolved, between *Allomyces* and humans.

In contrast to the anisogamous species, the sexual cycle in the isogamous species, *A. moniliformis* and *A. neomoniliformis*, has been less extensively studied. Isolates of these species produce morphologically identical, motile, colorless gametes of equal size that fuse to form pairs prior to zygote formation (Olson 1980); however, it is not known if pheromone signals or mating types are involved in driving the fusion between isogametes, and the molecular details of gamete interaction largely remain unclear.

Here, we present the analysis of transcriptomes of seven samples encompassing five isolates of *Allomyces*. Three strains represent the anisogamous mating system and the remaining two isolates are isogamous. We use comparative genomics of variants based on the transcripts from each strain to infer population relationships within the five strains. Further, we use these population relationships to analyze conserved gene expression between isogamous and anisogamous strains to understand the genome-wide changes associated with the evolution of anisogamy in *Allomyces*. We investigate gamete-type specific gene expression within and between the anisogamous strains and infer rates of evolution in male- and female-specific transcripts to understand the evolutionary constraints on these genes.

We find that anisogamous strains cluster together and away from isogamous strains, indicating that the transition to anisogamy likely happened once in *Allomyces*. Anisogamous strains expressed several unique transcripts including homologs of known sex determination genes in fungi and animals. Moreover, we found female-specific or male-specific expression patterns for several genes, some of which also showed signatures of rapid evolution. Interestingly, these included homologs of genes involved in Ca^2+^ ion transport, carotene biosynthesis, terpene metabolism and other sex-specific processes that are implicated as important contributors to gamete interaction in the anisogamous species of *Allomyces*. Overall, our analysis highlights conserved mechanism of gamete interactions in Ophisthokonts. We further report several candidate genes involved in this process and propose *Allomyces* as an ideal model system with which to study genetics underlying the evolution of gamete dimorphism in eukaryotes.

## Materials and Methods

### Strains and culturing conditions

The five strains used in this study are shown in Table 1. Three strains, Jun 1987, Neo and Burma 3-35, were purchased as filter paper stocks of dried sporangia from the Fungal Genetics Stock Center. Each of these strains was revived by immersing a piece of the filter paper in 5 mL of sterile water containing 4 boiled hemp seeds incubated at room temperature. Hydration releases motile zoospores from the sporangia that germinate to develop filamentous colonies. The plates were monitored daily for the presence of filamentous growth and a colony germinated from a single zoospore was transferred using glass microcapillary to YPSs agar (4g yeast extract, 15g soluble starch, 0.5g MgSO_4_ 5H_2_O, 1g K_2_HPO_4_, 20g agar, 1 L water, pH 7) containing 100 ug/mL ampicillin. Two strains, Allo1 and Allo2, were isolated from soil samples collected in Michigan and Virginia, respectively. Briefly, we collected soil from dried roadside ditches. A pinch of soil sample was added to 5 mL of sterile water containing four boiled hemp seeds. We incubated the plates at room temperature in condensation chambers and monitored daily for the presence of swimming zoospores and hemp-associated filamentous growth of *Allomyces*. A filamentous colony started from a single zoospore was isolated using glass microcapillary and washed serially with sterile water before transferring it to YPSs agar containing ampicillin as described before. All YPSs plates were incubated at room temperature. All cultures were maintained by transferring them to fresh medium every 15 days.

**Table 1.**
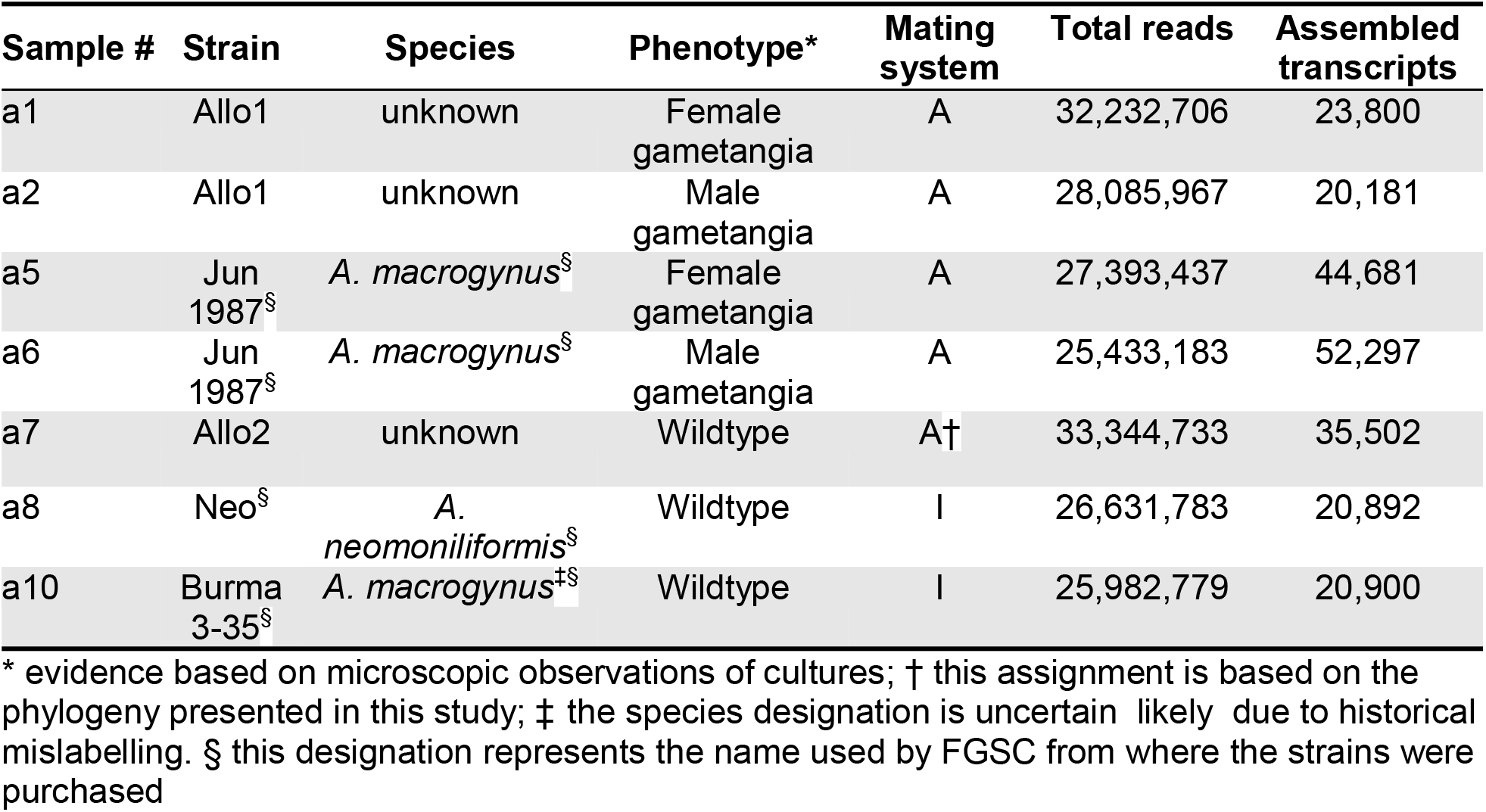
Samples used in this study. Transcriptomes from seven samples encompassing five strains were sequenced. Corresponding strain and species names are shown along with the details of life cycle stage and mating system. (A) refers to anisogamous and (I) refers to isogamous mating systems.

### RNA extraction, library preparation and transcriptome sequencing

We used 11-day old cultures for RNA extraction using Trizol reagent and Qiagen RNAeasy mini kit. We extracted total RNA from 7 samples as shown in Table 1. For samples a1, a2, a5 and a6, we extracted RNA from predominantly female (white gametangia) and predominantly male (orange gametangia) sectors of the colony (Figure 1; Table 1). Approximately 1 cm^2^ piece of the colony was used for RNA extraction per sample. For each strain, 50 μL of ~70-200 ng/ul total RNA, measured using bioanalyzer, was used for polyA library preparation without removing ribosomal RNA. The library preparation and sequencing was carried out at Yale Center for Genome Analysis (YCGA) using Illumina Hiseq 2500 paired-end high throughput 2X75 technology with insert sizes ranging from 130-200 bp. The raw reads have been submitted to NCBI under the accession #PRJNA421187.

**Figure 1.**
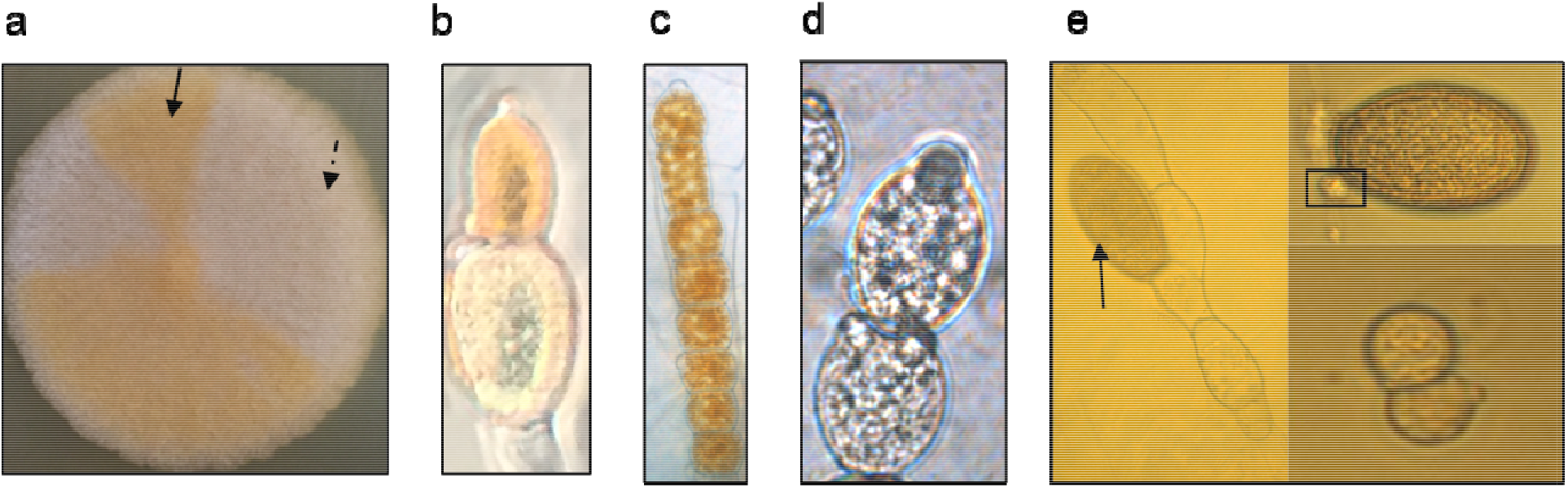
Sexual development in *Allomyces*. (a) A colony of an anisogamous strain is shown with orange sector that contains predominantly male (solid arrow) and white sector that contains predominantly female (dashed arrow) gametangia; (b) microscopic view of the hermaphroditic arrangement of gametangia in wildtype anisogamous strain; (c) microscopic view of a chain of predominantly male gametangia; (d) microscopic view of predominantly female gametangia; (e) microscopic view of isogamous development including a resting sporangium (arrow) representing meiotic stage, a meiospore (black rectangle) released from the resting sporangium and two encysted meiospores, each with a papillum (bud-like protrusion) from which isogametes are released.

### Sequence quality control and transcriptome assembly

Quality of the sequencing reads was verified using FastQC (http://www.bioinformatics.babraham.ac.uk/projects/fastqc/). We assembled unique transcriptomes for each sample using Trinity v2.2.0 (Grabherr et al. 2011) with the --normalize_max_read_cov 50 option. We ran Trimmomatic (Bolger, Lohse, and Usadel 2014) within Trinity using the --trimmomatic flag to remove k-mer, adapter contamination, and low quality nucleotides at the beginning and end of the reads. For this, we used default values including 2:30:10 SLIDINGWINDOW:4:5 LEADING:5 TRAILING:5 MINLEN:25. We also used the –jaccard_clip option to minimize the fusing of transcripts due to overlapping untranslated regions in the fungal genome. We obtained an average of ~20M reads for each of the seven samples (Table 1), yielding an average depth of ~50X per sample in *de novo* assemblies.

### Sequence alignment and filtering

The individual transcriptomes were aligned using ProgressiveCactus (Paten et al. 2011) with default alignment parameters on Arizona State University’s Ocotillo cluster. Using hal2maf included with ProgressiveCactus, we converted the binary hal file to a maf alignment file. A python script (mafTransToFasta) was used to extract aligned sequences from the maf file and output them in fasta format. We further subset this alignment with a python script (filterCDS.py), which extracted coding sequences by determining the most common open reading frame for each aligned block. We selected consensus open reading frames for each gene if they contained at least 200 nucleotides.

Using the subset coding sequences, we employed a series of python scripts to perform a variety of analyses on the coding sequence alignment. We used sampleMatrix.py to identify the total number of genes aligned between samples. We used subsetAlignment.py to extract all aligned blocks containing samples a1, a2, a5, and a6 as well as all alignment blocks containing only samples a1, a2, a5, and a6. This resulted in one file containing all aligned genes found in known anisogamous samples and one file with genes uniquely aligned between anisogamous samples. We similarly found genes that are uniquely expressed in isogamous samples a8 and a10. All programs for filtering and analyzing transcriptome alignments can be found at https://github.com/icwells/Quoll.

### Predicting gene function

We conducted a series of blast queries with NCBI BLAST+ (Camacho et al. 2009) on Arizona State University’s Ocotillo cluster. The nucleotide sequences were queried against the NCBI RefSeq database using blastn and against the SwissProt database using blastx (Madden 2013). We also used TransDecoder (https://transdecoder.github.io) on the nucleotide sequences to convert them into amino acids and used the output for blastp against the SwissProt database. We merged the blast output using the blastToCSV.py script and manually queried the NCBI protein database for full protein names. Additionally, we used the output from blastx to perform hmmscan (Finn, Clements, and Eddy 2011). The i-value and e-value cutoffs of 0.001 in the hmmscan output were used to predict gene function.

### Analysis of rate of evolution

To determine the rates of evolution for each set of orthologs, we used alignments between the 7 samples to estimate lineage-specific branch lengths. We assumed JC69 as the model of substitution and ran each alignment using baseml in paml 4.8 (Yang 1997) with parameters including clock=0, Mgene=0, fix_kappa=0, kappa =5, fix_alpha= 0, alpha = 0.1, Malpha=0, ncatG=5, RateAncestor=0, method=0, Small_Diff =1e-6 and cleandata=0.

### Phylogenetic analysis and ancestral character state reconstruction

We repeated the alignment process including the *Catenaria anguillulae* (NCBI accession # PRJNA330705) transcriptome as an outgroup. We identified homologs of 64 ORFs in each of the 7 *Allomyces* samples and *C. anguillulae*. These 64 ORFs were used to construct a phylogeny to infer the evolution of anisogamy in *Allomyces*. Briefly, we concatenated the 64 ORFs to create a ‘pseudo-genome” for each sample and for *C. anguillulae*. The pseudo-genomes of the seven samples and *C. anguillulae* were aligned using clustalw with default parameters (Thompson, Gibson, and Higgins 2002). Next, we constructed a maximum likelihood phylogeny with 1000 bootstraps using CLC 8.0 genomics workbench (https://www.qiagenbioinformatics.com/). We also constructed 10 alternate phylogenies using random subsets of the 64 ORFs to verify robustness of our species tree. All iterations supported the genetic relationships predicted in the phylogeny obtained using 64 ORFs. Furthermore, we constructed 64 individual gene trees and found that 97% of them (or 62 out of 64 trees) are consistent with the concatenated ‘species’ tree. Further, one of the two trees that were not consistent with the species tree, became consistent after choosing a different isoform. The phylogeny based on 64 ORFs was used to perform maximum likelihood ancestral character state reconstruction using Mk1 model in MESQUITE (Maddison and Maddison 2001).

## Results

### Anisogamy represents a derived state and likely evolved once in *Allomyces*

Theoretical studies have consistently predicted that anisogamy evolved from isogamous ancestors (Matsuda and Abrams 1999; Lehtonen, Kokko, and Parker 2016). The transition is justified because gamete dimorphism can increase fitness over monomorphism, either by optimizing mating encounters, reducing cytoplasmic conflict, or maximizing fertilization (Togashi and Cox 2011). Thus, it is expected that anisogamy is a derived state in taxa, such as *Allomyces*, in which closely related strains show either isogamous or anisogamous mating systems.

To determine whether anisogamy is mono- or polyphyletic and whether it represents a derived state in *Allomyces*, we investigated how the two mating systems are distributed across the phylogeny of our samples, rooted using a closely related isogamous species, *C. anguillulae* as an outgroup (Table 1; Figure 2). The anisogamous strains formed a well-supported clade separated from the isogamous samples, suggesting that anisogamy evolved once in *Allomyces*. Maximum likelihood ancestral character state reconstruction suggested that the ancestor of all *Allomyces* likely had isogamous mating system, which is maintained in two strains (Neo and Burma 3-35) included in our study (Figure 2). Thus, anisogamy likely represents a derived state in *Allomyces*, consistent with the theoretical predictions and reminiscent of the observations made previously in marine green algae (Wiese, Wiese, and Edwards 1979).

**Figure 2.**
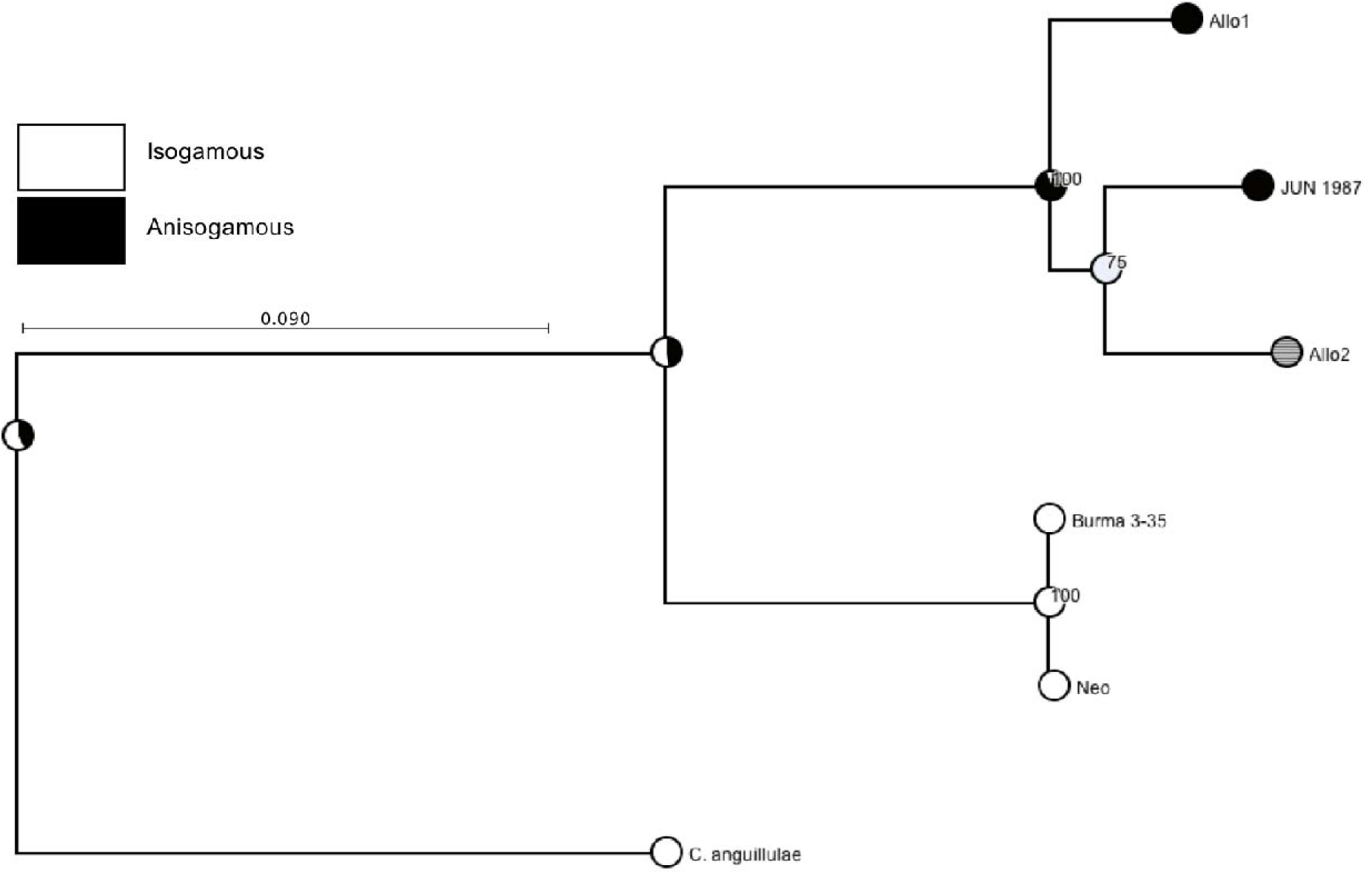
Evolution of anisogamy in *Allomyces*. Maximum likelihood phylogeny was constructed using the ‘pseudo-genomes’ (see methods for details) generated by concatenation of 64 orthologs shared between the samples a2, a6, a7, a8, and a10 and the outgroup (Supp. File S3). Bootstrap support is shown at internal nodes for 1000 replicates. The phylogeny was used in ancestral character state reconstruction in MESQUITE using the likelihood approach. Pie charts at the nodes represent the proportional likelihoods for each mating system given in the legend: isogamous (white); anisogamous (black); grey circles represent missing or unknown data. Alternative phylogenies including samples a1 and a5 are provided in the Supp. Figure S1.

Strain Allo2 is nested within the anisogamous species (Figure 2, Figure S1). We also found that this strain shares comparatively more transcripts with the anisogamous samples a1, a2, a5 or a6 than with the isogamous samples a8 and a10 in our study (Figure 3a, Figure S2d, Supp. File S1). These results suggest that Allo2 may be anisogamous; however, we have been unable to reproducibly induce the anisogamous hermaphroditic hyphae in the laboratory cultures of this strain. Further, the strain, Burma 3-35 was historically reported as being anisogamous (Olson and Nielsen 1981), but forms a well-supported clade in our analysis with the isogamous strain, Neo (Olson 1980) (Figure 1e, Figure 2, Figure S1). Burma 3-35 also shares most genes with the strain Neo (Figure 2a, Figure S2c) and has not shown anisogamous mating structures in laboratory cultures, even after repeated trials (data not shown). This suggests that either the historical strain identification, or the morphologically-based species concepts in *Allomyces*, need to be revised, as suggested previously (Porter et al. 2011). We obtained both, Burma 3-35 and Neo from the FGSC. If the strain Burma 3-35 indeed shows anisogamous reproduction in future studies, then our analysis would indicate that anisogamy may be polyphyletic and evolved at least twice independently in *Allomyces*. This issue will be revisited when more data on these and additional *Allomyces* strains become available.

**Figure 3.**
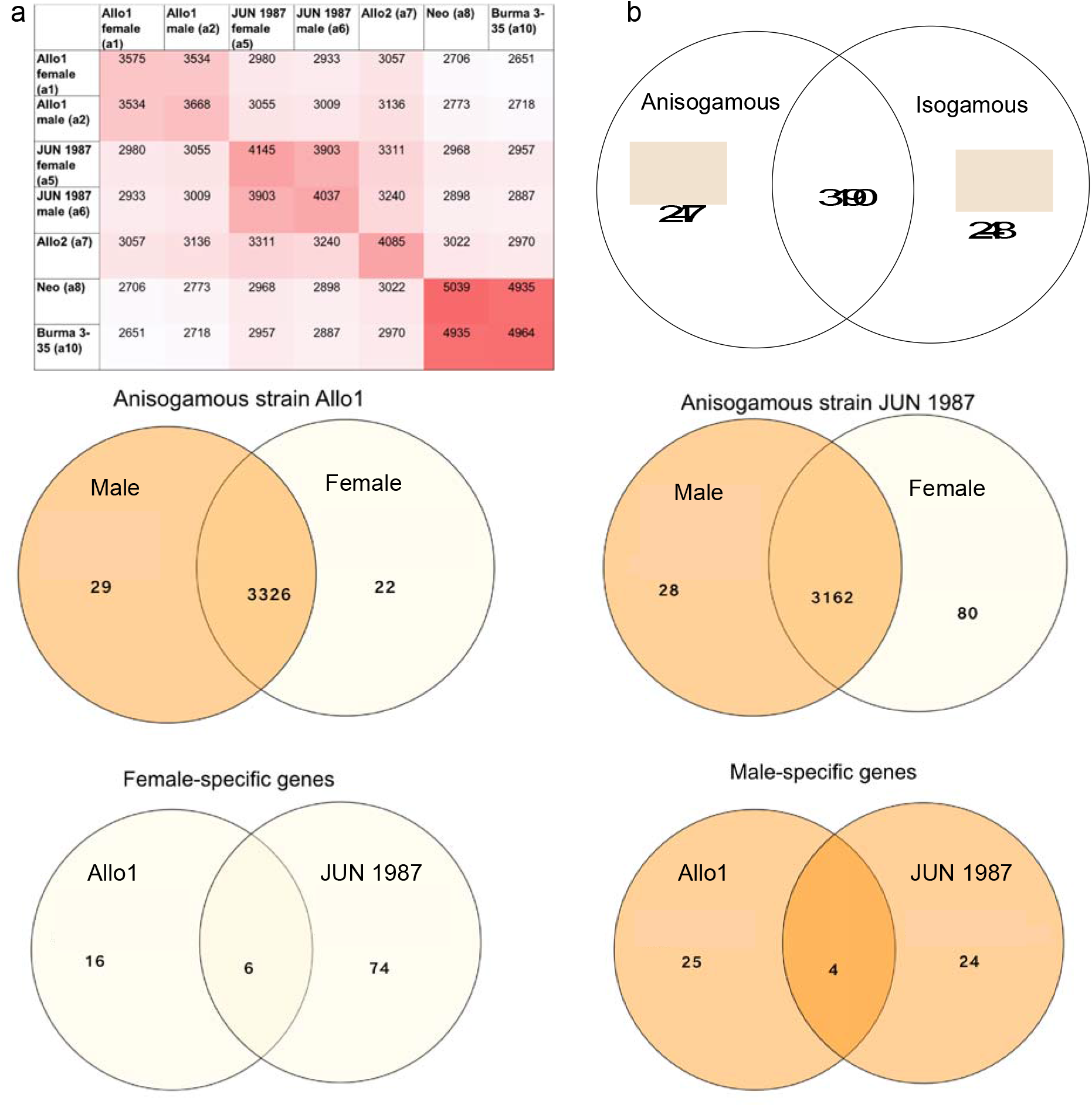
Overlap of assembled transcripts. The number of homologs shared between the 7 de novo assembled transcriptomes is shown. We inferred that a homolog existed if the sequence aligned between samples using Cactus (Paten et al. 2011). (a) a heatmap showing the number of homologs, including isoforms, shared between each pair of samples (b-f) venn diagrams are shown for the overlap between (b) isogamous samples a8 and a10 and anisogamous samples a1, a2, a5, and a6; (c) male and female samples of the anisogamous strain Allo1 (d) male and female samples of the anisogamous strain JUN 1987; (e) female-specific genes identified in c and d and (f) male-specific genes identified in c and d. For the venn diagrams, we did not distinguish between the isoforms of the same gene.

### Anisogamous strains show shared ancestral history and subsequent divergence from isogamous strains

The continuum of isogamous and anisogamous mating systems in closely related species offers an opportunity to investigate how genomes have been shaped following the transition to anisogamy in *Allomyces*. We compared the transcriptomes between anisogamous and isogamous samples to identify genes that are expressed in samples of both mating systems. Overall, the number of genes shared between pairs of samples corroborate phylogenetic relationships (Figure 3a). For instance, the two isogamous samples share more genes with each other than with either anisogamous sample (Figure 3a). However, there are many genes shared between the mating systems as well. Specifically, we found that anisogamous samples a1, a2, a5, or a6 express 3190 genes that are also expressed in either isogamous strains, a8 or a10 (Figure 3b, Supp. File S1), amounting to ~86% of all genes in our analysis. This reflects the recent genetic ancestry of the two mating systems represented in these five strains.

We also identified genes that are uniquely expressed in samples representing each mating system. We found 247 genes that are expressed in at least one of the anisogamous samples a1, a2, a5 or a6 but not in either of the isogamous samples a8 or a10 (Figure 3b). This number is further raised to 257 after including 10 genes that are uniquely expressed in the strain Allo2 (sample 7), which is likely anisogamous (Figure 2, Figure S2, Supp. File S1). The isogamous samples a8 and a10 expressed at least 229 genes that were not expressed in either of the anisogamous samples including the strain Allo2. If Allo2 is excluded from the analysis, there were 248 genes that were uniquely expressed in isogamous samples (Figure 3b). We note that with our limited sample size, these numbers and exact genes are not reflective of all possible evolutionary relationships between anisogamous and isogamous species, but given the divergent representation across mating systems we likely have identified major similarities and differences between them.

The genes that are uniquely expressed in each mating system likely reflect the diverse ecological selective pressures experienced by the lineages. We investigated the homology of the uniquely expressed genes against the known protein databases to understand their putative functions (Supp. File S2). We found that orthologs of a few genes involved in gamete-interaction, fertilization, and sexual development in fungi and animals are uniquely expressed in anisogamous samples (Supp. Files S1, S2). For instance, anisogamous samples expressed homologs of a gene containing the major sperm protein (MSP) domain that is important for sperm motility in a wide range of eukaryotic species (Tarr and Scott 2005; Tarr and Scott 2004). The MSP, first discovered in nematodes, is known to additionally act as an oocyte development stimulating hormone in the model hermaphroditic nematode *C. elegans* (Miller et al. 2001). Interestingly, we detected transcripts of the corresponding gene in both the predominantly female and predominantly male samples of the anisogamous strains (Supp. File S1). Also, anisogamous samples expressed orthologs of gelsolin-repeat containing protein and several genes involved in flagellar motility, which may represent candidate loci contributing to motility dimorphism in *Allomyces* (Pommerville 1978; Fritz-Laylin, Lord, and Mullins 2016). Other examples of genes uniquely expressed in anisogamous samples included an ortholog of Ste20, a gene regulating the mating process in many asco- and basidiomycete fungi (Dettmann et al. 2014), and orthologs of enzymes involved in mevalonate pathway and carotene biosynthesis, both of which are likely involved in male pigment production in the anisogamous species of *Allomyces* (Emerson and Fox 1940; Goldstein and Brown 1990). This category also included orthologs of genes coding for enzymes involved in sterol metabolism, which likely contributes to the synthesis of sirenin-the pheromone produced by *Allomyces* female gametes (Machlis 1958). Amongst genes that were uniquely expressed in isogamous samples, we found putative orthologs of several sperm-associated proteins; however, many of these produced only a weak match with the corresponding protein in the PFAM and TIGRFAM databases during the homology searches using hmmscan (Supp. File S2).

### Male and female sexes show strain-specific gene expression

We investigated sex-specific gene expression patterns in each of the two anisogamous strains, Allo1 (samples a1 and a2) and JUN 1987 (samples a5 and a6) (Figure 4). For each strain, we found that most genes were expressed in both sexes, with only a fraction of the genes uniquely expressed in either sex (Figure 3c and 3d). Moreover, the number of sex-specific genes varied between the two strains. Although the numbers were comparable between strains for the male-specific genes (29 in Allo1 and 28 in JUN 1987), there was greater discrepancy in the number of female-specific genes, with JUN 1987 females expressing around four times as many unique genes as Allo1 females (Figure 3c and 3d). This may be due to our use of transcriptome sequence instead of genome sequences. Another explanation is that samples were collected from predominantly, rather than exclusively, male or female sectors of the colony. If the ratio of female:male gametangia in the predominantly female samples varied between the strains, then we expect that difference to be reflected in the number of sex-specific genes in each strain. We further asked if the same genes show sex-specific expression in the anisogamous strains. We found very little overlap between the strains, which shared only a few female-specific (Figure 3e) and male-specific (Figure 3f) genes. This suggests that gene expression in our study was more strongly influenced by strain-specific factors rather than sex. Supporting this notion, we found stronger correlation in gene expression between sexes within a strain (Figure 4A and 4B) as compared to the correlation between strains for either sex (Figure 4C and 4D). Analysis of differential gene expression between exclusively-female and exclusively-male mutants of each strain will be necessary to identify genes showing biased expression in either sex. Such genetic mutants of *Allomyces* have been described in the literature and can be induced using X-ray and chemical mutagenesis (Rønne and Olson 1976; Ojha and Turian 1978). Alternatively, the limited overlap of sex-specific genes between strains might indicate that anisogamy evolved independently in strains Allo1 and JUN 1987; however, our analysis indicates that this is an unlikely scenario (Figure 2).

**Figure 4.**
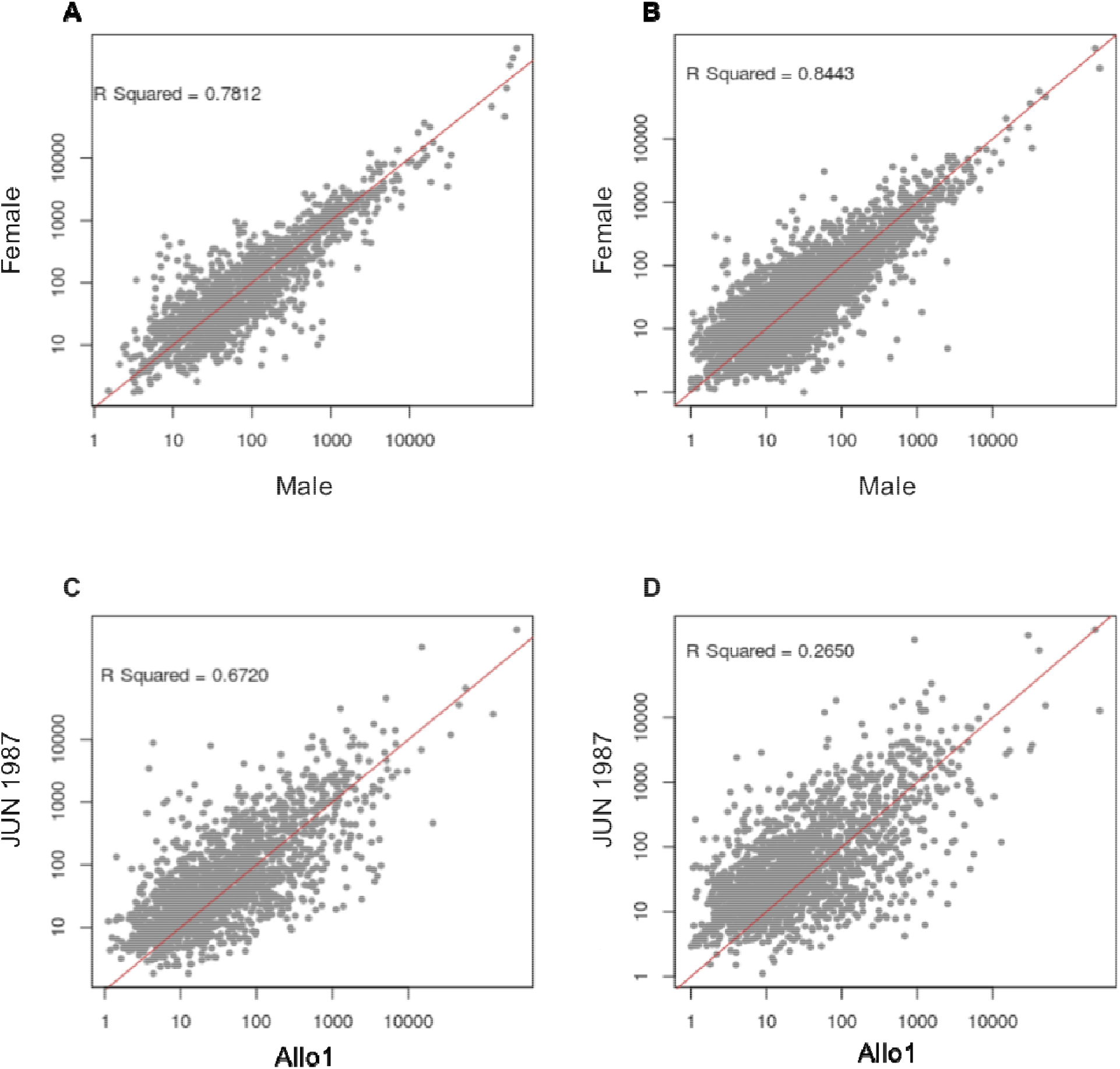
Gene expression in males and females. A strong correlation between the sexes is seen for the anisogamous strains Allo1 (A) and JUN 1987 (B). Gene expression is less correlated between the strains for either male (C) or female (D) sex. The grey dots in all plots represent FPKM values for individual genes.

### Many genes show mating system-specific and lineage-specific patterns of accelerated evolution

Multiple studies have shown that sex-related genes show signatures of positive selection and rapid evolution (Swanson and Vacquier 2002; Ferris et al. 1997; Haerty et al. 2007; Papa et al. 2017). This phenomenon has been attributed to sexual conflict and selection, and more specifically to the evolutionary arms race between sexes, which can continuously select for new variants at the respective loci (Clark, Aagaard, and Swanson 2006; Fiumera, Dumont, and Clark 2005; Gavrilets 2000). We surveyed the *Allomyces* transcriptomes for genes that show high numbers of substitutions between the lineages used in our analysis. If there are strong allele-specific expression effects, we may observe fewer substitutions, but we do not expect this to inflate our estimates. Additionally, because we lacked information on optimum open reading frames, we could not use Ka/Ks ratio test to infer non-neutral patterns of evolution. Instead, we estimated lengths of individual gene trees using total substitution rates and used them to infer mating system-specific and lineage-specific patterns of evolution at these loci (Figure 5). We found that many genes showed high levels of sequence divergence between mating systems, likely indicating saturation of nucleotide changes (Supp. File S5). Moreover, lineage-specific patterns of substitutions resulted in the respective lineages having extremely long branches in gene trees (Supp Table S4). The branch lengths were robust to substitution models and model parameters assumed in our analysis and instead stem from the patterns of substitutions between samples (Supp. File S4 and Supp. File S5). It is likely that the unexpectedly high levels of sequence divergence reflect gene duplication and subsequent diversification for the respective genes (Heger and Ponting 2007; Obbard et al. 2006). Alternatively, it may indicate rapid evolution, especially of the genes that show multiple polymorphisms between male and female samples of a given lineage.

**Figure 5.**
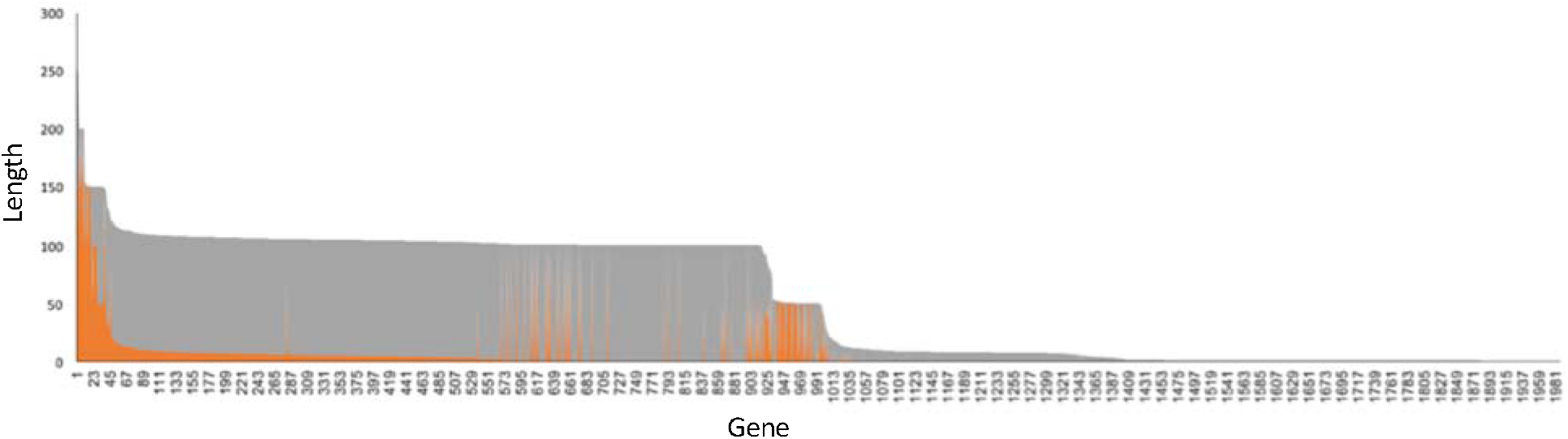
High levels of sequence divergence and genetic saturation. The distribution of tree lengths (y-axis) is shown for genes represented as unique numbers (x-axis). Sequences corresponding to the unique numbers along with the lineage-specific branch lengths are listed in Supp. File S5 and Supp. File S4, respectively. The anisogamous lineages (orange bars) including samples a1, a2, a5, a6 and a7 together show lower tree lengths than the isogamous lineages (grey bars) including samples a8 and a10.

The signs of rapid evolution in many genes may reflect various selective pressures including ecological and sexual selection experienced by each lineage analyzed in our transcriptomes. We used hmmscan to identify putative function of the genes showing long branches (Supp. File S4). Some of the genes that are expected to contribute to sexual dimorphism in *Allomyces* showed high levels of substitutions in a lineage-specific manner. For instance, we found the putative homologs of the gene encoding diphosphomevalonate decarboxylase-an enzyme involved in mevalonate pathway, to be rapidly evolving in the anisogamous lineages but not in the isogamous lineages (Supp. File S5, Supp. File S4). The mevalonate pathway likely contributes to two gamete-specific traits in the anisogamous species of *Allomyces*. First, it may provide precursor products required for the synthesis of the carotenoid pigments specific to the male gametes (Goldstein and Brown 1990). Second, it may be involved in the synthesis of sirenin-a pheromone that is secreted by the female gametes (Goldstein and Brown 1990). Sirenin is a sesquiterpene molecule known to mimic the action of mammalian steroid hormone, progesterone (Syeda et al. 2016), whose synthesis requires the mevalonate pathway in animals (Hellig and Savard 1965; Tuckey 2005). Similar lineage-specific patterns of substitutions were apparent for other genes involved in the mevalonate pathway including putative homologs of HMG CoA synthetase and 3-oxoacyl synthase (Supp. File S4).

We found that a gene that was identified as a putative homolog of sex-lethal family splicing factor showed long branches in isogamous but not in anisogamous lineages. Similar patterns were observed for the homolog of the gene encoding lycopene synthase-an enzyme likely involved in carotenoid biosynthesis in the male gametes of *Allomyces* (Emerson and Fox 1940) and for the homologs of *CatSper* genes, which show sperm-specific expression in metazoa (Strünker et al. 2011). It is not known whether any of these genes show signatures of positive selection. Currently, efforts are underway to predict open reading frames for the genes identified in our transcriptomes. The resulting data along with functional genomics studies will be necessary to advance our understanding of the sex determinants and the evolutionary constraints under which they evolve in *Allomyces*.

## Discussion

Sexual reproduction in eukaryotes repeatedly evolved from isogamy to anisogamy. The first molecular genetic evidence for this transition was presented by Nozaki et al. (2006) in a group of marine algae. Since then, subsequent studies have continued to reveal genetic rewiring responsible for the evolution of anisogamy within the algal lineages. Unfortunately, similar molecular insights remain elusive for most other groups of eukaryotes. Here we introduce *Allomyces* - an early diverging fungus that provides an excellent empirical model to understand the evolution of anisogamy in Ophisthokonts-the clade shared between fungi and animals. *Allomyces* shows a continuum of the two mating systems in closely related species (Emerson 1941; Porter et al. 2011), allowing the study of genetic relationships between anisogametes and their isogamous ancestors. Using transcriptome analyses of five isolates of *Allomyces*, we find support for the hypothesis that anisogamy evolved once in *Allomyces* and that the ancestor of all *Allomyces* was isogamous.

Our phylogenetic analysis is consistent with the morphospecies-based classification proposed for *Allomyces* in which strains with distinct mating systems occupy distinct clades. Previously, Porter et al. (2011) constructed rDNA-based phylogeny for additional strains and found it to be incongruent with the morphospecies based classification, suggesting that the species concepts or strain identification in *Allomyces* may need to be revised (Porter et al. 2011). Corroborating their observation, our analysis indicates that at least one of the strains used in our study is likely to be historically mislabelled. The strain Burma 3-35 or ATCC38327 is originally documented to belong to the model anisogamous species *A. macrogynus* (Olson and Nielsen 1981) whose genome sequence is available (NC_001715). Although the sequenced strain is thought to be Burma 3-35 (https://www.ncbi.nlm.nih.gov/genome/?term=txid28583 [Organism:exp]), we observed that the strain JUN 1987 shows almost perfect similarity to the sequenced genome (NC_001715). In contrast, Burma 3-35 shared more transcripts and higher genetic similarity with the isogamous strain Neo used in our study than with any other strain. Taken together, we propose that further genetic verification is imminent to identify the original Burma 3-35 strain of the model species *A. macrogynus*. Additionally, our study did not include isolates representing the morphospecies *Brachy-Allomyces* (Olson 1984). Hence, a detailed molecular phylogenetic analysis inclusive of such isolates is necessary to verify whether species with distinct mating systems indeed occupy distinct clades in *Allomyces*.

We found that isogamous and anisogamous strains shared many genes. In fact ~80% (or 2862) of the genes identified in our de novo transcriptomes showed expression in all seven samples, and only 16 showed expression specific to a single sample (Table S1). The reference genome of the model anisogamous species, *A. macrogynus* (NC_001715) is estimated to contain ~19,500 protein-coding genes. Hence, our analysis may be limited to ~16% of the functional genome, if all species included in our study have comparable gene content.

Interestingly, many transcripts produced significant matches with conserved domains found in genes involved in gamete interaction and sex determination in other Ophisthokonts, which allowed us to identify several putative sex determinants in both isogamous and anisogamous samples (Supp. File S2). The putative sex determinants roughly fall into two categories including genes known to show sex-biased expression in metazoa and genes involved in fungal mating processes. In the first category are homologs of sex-lethal family splicing factor, which induces female-specific patterns of alternative pre-mRNA splicing and triggers female sexual differentiation in *Drosophila* (Penalva and Sánchez 2003; Wang, Dong, and Bell 1997). We also found orthologs of male-specific genes including those containing the gelsolin repeat containing domain, motile sperm (MSP) domain, spermine synthase domain, NYD-SP28 sperm tail domain, speriolin N terminus domain, the spermatogenesis-associated protein domains (SRSP1 and SPATA19), sperm acrosome-associated protein (SPACA7) domain and the TEX domains, which are found in genes showing testis-specific expression (Wu et al. 2003, 14). In addition, we identified multiple orthologs of CATSPER genes, which encode sperm-specific Ca^2+^ channels that regulate influx of calcium into mammalian sperm during hyperactivation (Strünker et al. 2011). Genes encoding *CatSper* channels have been previously reported in the model anisogamous species *A. macrogynus* (Cai and Clapham 2012). Also, it has been shown that Ca^2+^ signaling plays an important regulatory role in gamete fusion in the anisogamous species of *Allomyces* (Pommerville 1977; Pommerville, Strickland, and Harding 1990). These discoveries along with our finding that homologs of CATSPERB, CATSPERG and CATSPERD are expressed during the sexual stage in *Allomyces* indicate that gamete fusion in anisogamous species of *Allomyces* likely involves Ca^2^+ signaling through *CatSper* channels, similar to that in mammals. We were unable to perform the analysis of differential expression due to a limited number of biological replicates; however, efforts are currently underway to determine if *CatSper* genes show sex-specific expression in the anisogamous species of *Allomyces*. Male-biased expression of *CatSper* in a basal fungus like *Allomyces* will raise the hypothesis that the role of *CatSper* as sex-specific Ca^2^+ channel evolved in the ancestor of all Ophisthokonts-the clade shared between animals and fungi. Alternatively, it could represent convergent evolution.

There is direct evidence suggesting that gamete interactions in the anisogamous species of *Allomyces* involve the same molecular events as in the egg-sperm interaction in humans. Specifically, Syeda et al (2016) discovered that sirenin – a pheromone secreted by the female gametes of *A. macrogynus* – mimics the action of progesterone and hyperactivates human sperm *in vitro* (Syeda et al. 2016). Sirenin is a sesquiterpene molecule that is known to be secreted by female gametes and attracts conspecific and heterospecific male gametes in *Allomyces* (Machlis, Nutting, and Rapoport 1968; Machlis 1968). It is unknown what pathways are responsible for the synthesis of sirenin in *Allomyces;* however, the process likely involves terpenoid precursors (Machlis, Nutting, and Rapoport 1968). Interestingly, these precursors, which are created by the mevalonate pathway, are important intermediates from which all sterols and steroid-based hormones are produced in animals (Goldstein and Brown 1990). Our transcriptome analysis revealed that *Allomyces* expresses orthologs of several genes involved in the mevalonate/HMG-CoA reductase pathway including mevalonate kinase, diphosphomevalonate decarboxylase, phosphomevalonate kinase, and HMG-CoA synthase (Spp. Table S2). In addition, we found that isogamous and anisogamous samples expressed orthologs of genes involved in progesterone biosynthesis, metabolism and signal processing including genes encoding terpene synthase, 3-beta hydroxysteroid dehydrogenase, an oxysterol binding protein, 3-oxo-5-alpha-steroid 4-dehydrogenase, sterol-sensing domain of SREBP cleavage-activating protein, hydroxy-steroid dehydrogenase and the progesterone receptor (Supp Table S2). Some of these genes showed female-specific expression patterns in our samples (Supp. File S1); however, quantitative evidence using differential expression analysis will be necessary to verify sex-biased expression of these genes and to further understand their role in sex determination in *Allomyces*. Although molecular genetics and functional genomics studies are essential to determine whether and how these genes regulate sexual differentiation in *Allomyces*, some of these genes showed signatures of rapid evolution (Supp. File S4), reminiscent of sex-related genes in other many other species (Swanson and Vacquier 2002), lending further support to the notion that the molecular machinery involved in sexual differentiation and gamete interaction may be conserved between *Allomyces* and animals including humans.

The second category of putative sex determinants that were detected in our transcriptomes include known fungal mating genes. Amongst these, we found orthologs of MFαprecursor, which undergoes proteolytic processing to produce mature *a* factor pheromone in *Saccharomyces cerevisiae* (Caplan et al. 1991). We also found expression of orthologs of Prm10P, a pheromone-regulated 5-transmembrane protein that is predicted to be involved in yeast mating (Bardwell 2005). In addition, we identified genes containing the sterile motif/pointed (SAM_PNT) domain, which is present in many proteins involved in yeast sexual differentiation (Kim and Bowie 2003). Amongst genes that are active in the mating process of non-yeast filamentous fungi, we could identify orthologs of velvet family genes, which are known to perform diverse roles in fungal morphogenesis and development including regulation of microconidia-macroconidia ratios in the heterothallic fungus *Fusarium verticilloides* (Li et al. 2006). We also found transcripts of a gene containing the AαY_MDB domain, which is responsible for mating-type dependent binding of Y protein to the AαZ protein of another mating type in the basidiomycete fungus *Schizophyllum commune* (Robertson et al. 1996).

In all fungi including *Allomyces*, the onset of sexual stage is defined by the onset of meiosis (Mirzaghaderi and Hörandl 2016). Not surprisingly, our transcriptomes included the meiosis-specific genes *Spo22, Rad21, Rad51, Dmc1*, and MAP kinases including *Ste20*, which are primarily expressed during sexual stage in the life cycles of many eukaryotes including fungi (Schurko and Logsdon 2008; Malik et al. 2008; Butler 2010; Dettmann et al. 2014). Lastly, we found orthologs of genes that are likely involved in carotene biosynthesis, which is regulated by environmental triggers including sexual stimulation in some fungi (Govind and Cerda-Olmedo 1986). For instance, the model fungus *Phycomyces blakesleeanus* in the subphylum mucormycotina derives carotenoid compounds from HMG-COA in the mevalonate pathway in response to sexual interactions (Polaino et al. 2012, 2010; Govind and Cerda-Olmedo 1986). Genes responsible for carotenogenesis in these fungi include those encoding lycopene cyclase, phytoene desaturase, squalene synthase, and squalene-associated FAD-dependent desaturase, all of which were expressed in our transcriptome samples and showed lineage-specific signatures of rapid evolution (Supp. File S4). The later two enzymes are also essential components of the pathway leading to biosynthesis of hopanoids (Sáenz et al. 2015), which are pentacyclic steroid-like compounds derived from squalene-a diterpene that is metabolized to produce several carotenoid compounds (Kannenberg and Poralla 1999; Rohmer, Bouvier, and Ourisson 1979). Expression of orthologs of these genes in *Allomyces* likely represents candidate loci contributing to the synthesis of orange pigment in male gametes. Emerson (1940) purified this pigment and found it to contain *γ* –carotene (Emerson and Fox 1940). Moreover, the female gametes in anisogamous species and the gametes of isogamous species seem to lack this pigment (Olson 1980; Olson 1984; Emerson 1941; Ji and Dayal 1971; McCranie 1942). Thus, the male gamete pigmentation represents a truly dimorphic character and the respective loci provide first examples of specific genomic regions associated with gamete dimorphism in *Allomyces*.

It remains unclear how many loci contribute to determine gamete size dimorphism in *Allomyces*. The gametes in the isogamous species are ~5 um in diameter, similar in size to the male gametes, and about half the size of the female gametes in the anisogamous species (Pommerville 1982; Pommerville and Fuller 1976; Olson 1984). Population genetic models have predicted that anisogamy can evolve when a single locus determining gamete size becomes linked to the sex determination locus (Charlesworth 1978). Although such simple genetic mechanisms are perhaps rare (Beukeboom and Perrin 2014), anisogamy may initially evolve from relatively few sex determination loci as found in marine algae in which small male gametes evolved from one of the isogametic mating types (Nozaki et al. 2006). Initial differences stemming from a few loci may quickly become reinforced leading to sexual dimorphism in traits other than gamete size. In many species, size dimorphism is usually accompanied by dimorphism in gamete motility (Hoekstra, Janz, and Schilstra 1984; Hoekstra 1984; Cox and Sethian 1985). In anisogamous species of *Allomyces*, dimorphism is apparent in gamete size, motility, pigmentation and pheromone production and likely involves additional molecular differences between male and female gametes. In terms of motility, the anisogamous system of *Allomyces* is similar to that of the green algae *Chlamyomonas* because both male and female gametes are motile (Pommerville 1978). However, unlike *C. reinhardtii* in which the two gamete types are comparably motile (Starr, Marner, and Jaenicke 1995; Lefebvre 1995), the female gametes of *Allomyces* move more sluggishly than the male gametes (Cox and Sethian 1985). The anisogamous system in *Allomyces* may represent an intermediate step towards the evolution of oogamy in which one gamete type (usually female) completely lacks flagellar apparatus and the associated motility (Kirk 2006). Indeed, the transition from isogamy to anisogamy (size dimorphism) to oogamy (size and motility differences) has been theoretically predicted (Wiese, Wiese, and Edwards 1979; Birky 1995) and empirically documented in the green algae (Nozaki et al. 2014). Oogamy is also the norm in metazoa (Kirk 2006; Leonard and Cordoba-Aguilar 2010). Analysis of mating systems in additional isolates of *Allomyces* will be necessary to determine whether oogamy has evolved in this genus.

The size dimorphism in anisogamy has been interpreted in game theoretical models as an unequal investment per gamete by the sexes (Bulmer and Parker 2002). Usually, male gametes (sperm) are smaller than female gametes (eggs). Accordingly, when a small sperm fuses with a relatively large egg to form a zygote, the female would have invested more resources per zygote than the male. In game-theory, the differences between the gamete size (investment) and number can form the primary distinction between the sexes, setting the stage for sexual conflict and selection in anisogamous systems (Smith 1977; Arnold and Duvall 1994). It is not clear whether loci contributing to gamete differentiation and sexual development in *Allomyces* experience sexual selection but given that we found signatures of rapid evolution in some of these genes, empirical analysis of mate competition is warranted. Our study shows that *Allomyces*, if developed further as a model in functional genomics, will offer exciting opportunities to explore the relationship between anisogamous mating, sexual selection and the evolution of sexual dimorphism.

*Allomyces* can also lend itself to experimentally test some of the theoretical models explaining the adaptive significance of anisogamy. For instance, the evolution of anisogamy is famously predicted to be a response to disruptive selection against intermediate-size gametes such that larger gametes ensure increased zygote survival and smaller gametes ensure greater fertilization success (Parker, Baker, and Smith 1972). Although anisogamy in *Allomyces* is thought to exemplify the assumptions and predictions of the PBS theory (Togashi and Cox 2011), these have never been tested empirically in *Allomyces*. First, it is not known whether there is a negative correlation between the size and number of gametes in *Allomyces*, which can set the stage for the evolution of anisogamy. Variation in gamete size in response to oxygen concentration has been documented (Olson and Rønne 1975) but it is unclear if it correlates with gamete number. Second, there is no experimental evidence for gamete competition in *Allomyces*, which is another requirement of PBS for species with external fertilization including *Allomyces* (Parker, Baker, and Smith 1972). Extensions of PBS using game theoretical approaches have shown that in the absence of gamete competition, sexual selection in the form of sperm competition may select for smaller size of sperm, especially in species with internal fertilization (Bulmer and Parker 2002). Although *Allomyces* has external fertilization, male-male competition could contribute to the maintenance, if not origin, of gamete dimorphism in the anisogamous species.

An alternative explanation of why anisogamy is prevalent is offered by the sperm limitation theory, which proposes that the increase in female gamete size is an adaptation to maximize fertilization success when mates are limited (Levitan 1993). This is an unlikely scenario in *Allomyces* because 1) gametes are formed on hermaphroditic hyphae facilitating gamete encounter (Pommerville 1982), 2) gametes are released synchronously facilitating contact (Sewall and Pommerville 1987; Morrison 1977) and 3) occasionally zygotes can form parthenogenetically allowing reproduction even in the absence of mates (Olson 1984). Lastly, according to the theory of uniparental inheritance, cytoplasmic elements may select for large gametes through competition for space and in turn select for nuclear genes that trade gamete size for number to maximize mating success (Murlas Cosmides and Tooby 1981; Hurst and Hamilton 1992). Eventually, a single allele for size is fixed, simply establishing the PBS scenario (Hoekstra 1987). The same theory also describes anisogamy as a response to selection against biparental inheritance of the cytoplasm, which can lower fitness of the zygotes if the parents carry incompatible cytoplasmic elements (Hurst and Hamilton 1992). Thus, anisogamy may be adaptive because it imposes uniparental inheritance by eliminating cytoplasm from the smaller gamete. In *Allomyces*, zygotes appear to retain paternal mitochondria, at least in the interspecies crosses (Borkhardt and Olson 1983). Also, parthenogenesis may eliminate constraints of intracellular conflict in *Allomyces* (Olson 1984). This suggests that uniparental organelle inheritance may not have caused the evolution of anisogamy in *Allomyces*, although it may be responsible for the maintenance of dimorphism once it was evolved.

It is not a trivial task to experimentally test the adaptive mechanisms underlying the evolution of anisogamy as proposed in the theoretical models. The conventional unicellular systems such as yeast are likely to prove suitable for studying the evolution of only a subset of dimorphic traits, for instance, size but not motility (Kawecki et al. 2012). Recent studies on the green algae have emphasized the significance of studying gamete dimorphism in non-traditional microbial models but thus far, they are the only biological systems available for empirical investigation into the molecular basis of anisogamy. Prior to the availability of genome sequence, decades of research on *Allomyces* has contributed several molecular and physiological details about its anisogamous life cycle. Our study upholds these efforts to show that *Allomyces* is likely to provide important insights into the evolutionary forces and genetic changes underlying sexual dimorphism at the level of gametes. We predict that future research on mating systems in *Allomyces* will have significant implications for understanding the evolution of sexual reproduction, and the diversification of Ophisthokonts.

## Acknowledgements

This work was supported by the startup funds to MAWS from the School of Life Sciences and Biodesign Institute, Arizona State University and by Systematics Research Fund, Royal Society of London to SSP. Christopher Castaldi and Erica Ryke, Yale University assisted with library preparation and sequencing. SSP would like to thank Dr. Joseph Heitman, Duke University and Dr. Tim James, University of Michigan for introducing her to *Allomyces* and for the initial guidance with culturing strains, respectively. We are grateful to Dr. Rebecca Zufall, University of Houston and Dr. Vikram Shende, California Institute for Biomedical Research for helpful discussions and comments on the manuscript.

